# Recursive Convolutional Neural Networks for Epigenomics

**DOI:** 10.1101/2020.04.02.021519

**Authors:** Aikaterini Symeonidi, Anguelos Nicolaou, Frank Johannes, Vincent Christlein

**Affiliations:** Technical University of Munich, Liesel-Beckmann-Str. 2, 85354 Freising, Germany; Friedrich-Alexander University Erlangen-Nürnberg, Martensstr. 3, 91058 Erlangen, Germany

**Author notes:** Equal contribution (Aikaterini Symeonidi, Anguelos Nicolaou).

## Abstract

Deep learning methods have proved to be powerful classification tools in the fields of structural and functional genomics. In this paper, we introduce a Recursive Convolutional Neural Networks (RCNN) for the analysis of epigenomic data. We focus on the task of predicting gene expression from the intensity of histone modifications. The proposed RCNN architecture can be applied to data of an arbitrary size, and has a single meta-parameter that quantifies the models capacity, thus making it flexible for experimenting. The proposed architecture outperforms state-of-the-art systems, while having several orders of magnitude fewer parameters.

## I. Introduction

All somatic cells of eukaryotic organisms contain the same DNA sequence, yet there are vast functional differences between cells across tissue types. It has become increasingly clear that these functional differences are partly determined by epigenetic factors that alter gene expression patterns without changing the underlying DNA sequence [1], [2]. Two key epigenetic mechanisms are DNA methylation [3] and modifications of histone proteins [4], [5], [6]. Histone modifications, are chemical modifications of the histone proteins around which the DNA is wrapped and have an indirect influence on gene expression by changing the accessibility of the DNA [7], [5]. The distribution of these epigenetic marks along the genome acts like a regulating mechanism that defines the functional state of a cell at a given time. As an example, it has been shown that circadian programming of cells, is realised through such epigenomic mechanisms, more specifically H3K27me3 and H3K4me3 regulate core circadian genes[8]. Numerous plant and animal studies have collected epigenomic data from different tissue types, developmental stages, disease states and environmental treatments [9], [10], [1], and have shown that this information is instrumental in elucidating key functional changes during cellular differentiation, disease pathology and for annotating causal mutations from genome-wide association (GWAS) mapping results [9], [11], [12]. To date, more than 100 histone modifications have been described [1], [4], [7], [13]. The large number of marks has led to the notion of a “histone code”, a layer of epigenetic information that is encoded by the combinatorial presence/absence patterns of specific histone modifications along the genome [6], [14]. Indeed, molecular studies have shown that histone modifications often act cooperatively in recruiting various chromatin readers and writers as well as transcriptional activators to particular genomic regions [15], [16], indicating that their combinatorial presence is required for proper gene regulation. Recent advances in next-generation sequencing (NGS) technologies now allow scientists to profile gene expression data along with histone modifications and other genomic and epigenomic signals at an unprecedented resolution and scale. The signal obtained from NGS technologies is a discrete signal over each position of the genome. The accumulation of such big datasets in the public domain provides novel opportunities for data mining.

Several computational models have been proposed, using regression models [17], [18], support vector machines [19], random forests [20], or rule-based models [21], for the task of predicting gene expression from histone modifications. However, the above approaches do not capture the combinatorial effect of histone modifications and require a great deal of effort in order to optimize the various model parameters. In recent developments it has been shown that deep learning trumps in performance aforementioned approaches[22], [23].

The principal contributions of this paper are: The introduction of recursive networks for epigenomic data analysis. The restriction of free parameters bellow the cardinality of of the train-set making over-fitting practically impossible, and large-scale cross-domain experimentation demonstrating the proposed models tend to model generic biological phenomena rather than dataset specific correlations.

## II. Related Work

### A. Deep Epigenomic Analysis

Neural networks have already been used for the task of gene expression prediction from histone modification marks. Singh et. al [22] proposed DeepChrome, a classical Convolutional Neural Network (CNN), with one convolutional layer and two fully connected layers. Dropout was employed to reduce over-fitting to the training data. The network consists in total of four layers and approximately 650 thousand free parameters and outperformed other machine learning approaches. Follow on work by the same group, proposed a second architecture, AttentiveChrome[23], a Recurrent Neural Network (RNN) with two levels of soft attention. This network also consists of four layers and reduces the free parameters to approximately 55 thousands. These models were applied to a collection of 56 human datasets each one containg the histone marks and the gene-expression for a specific cell-line. DeepChrome achieved an average AUC score 80.1%[23] and AttentiveChrome an average AUC score of 81.3% for the same datasets. We consider the experimental process a de-facto standard for comparing models that predict gene expression from histone marks. More details about the employed data and their analysis can be seen in IV-A. In Fig.1 examples of both expressed and not expressed samples as employed in DeepChrome and AttentiveChrome can be seen.

**Fig. 1.**
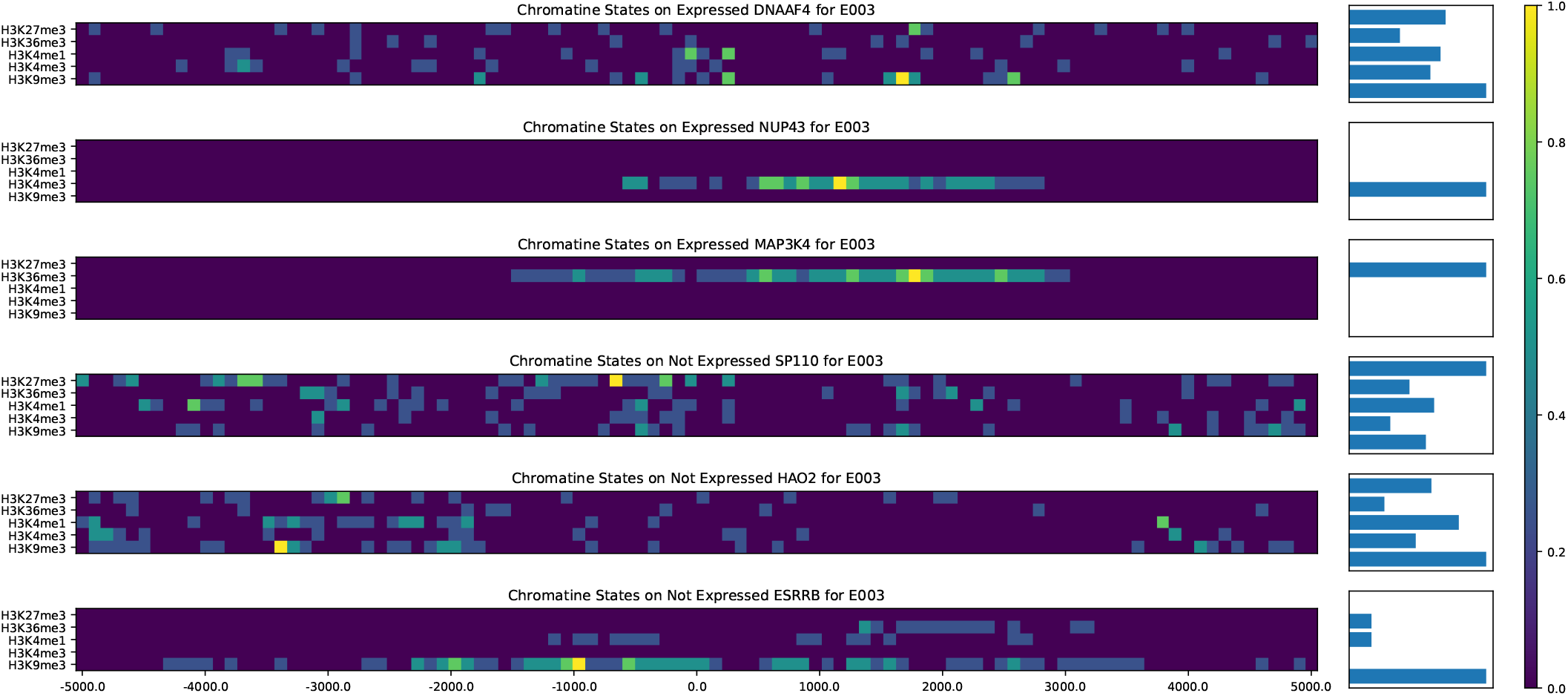
Chromatine states around the Transcription Start Site (TSS) of expressed and non-expressed genes. Barplots contain the sum per channel.

### B. Recursive Networks

Recursive Networks and Back-Propagation Through Structure (BPTS) were proposed in 1996 by Goller et al. [24] as a generalization of Back-Propagation Through Time (BPTT). They were introduced as a means of modeling information structured as directed acyclic graphs. The similarities between recursive and recurrent neural networks are very large and have occasionally been confused in older literature, since both have the acronym RNN. Most importantly, they both suffer from vanishing and exploding gradients [25]. In 2011, recursive networks were used for scene and language parsing [26] and achieved state-of-the art performance for those tasks. The model was not directly producing a global tree-structure; the optimal structure was estimated at inference-time with a greedy algorithm. Convolutional variants of recursive networks were first introduced by Socher et.al [27] for 3D object detection. Recursive convolutional networks allow for a fixed tree-structure and thus remove the need for a greedy algorithm at inference-time. Kim et. al [28]. employ a recursive CNN to the problem of image super-resolution. They propose to apply intermediate losses to each recursion layer, similar to the inception network [29] in order to alleviate vanishing/exploding gradients. Additionally, they employ skip connections from the input to the output which is beneficial for super-resolution.

## III. Methodology

We model the problem as one of sequence classification similar to other approaches[22], [23]. Specifically, we model the input as a five channel continuous 1D signal, and gene expression as a two-class problem: *“is expressed”* or *“is not expressed”*. The input signal is a quantification of histone modification marks on the genome, which is obtained by mapping snippets of these marks on the genome and counting their amount. Similarly, the expression signal is a quantification of RNA and is obtained by counting snippets on the genes of the genome and normalizing for factors like gene length and depth of sequencing. The genome samples consist of a couple of thousands of genes; depending on the data partitioning, this number can reach ten thousand examples while the input features are orders of magnitude larger. Therefore, one of the main problems is overfitting, which commonly occurs for models using genomic data or other biological data of equivalent cardinalities such as epigenomics. Romero et. Al [30] proposed that the ratio of number of inputs to the number of parameters is alarming and must be countered by reducing the number of free model parameters as established regularization techniques such as dropout are probably inefficient. Therefore, they proposed architectures that were primarily designed to drastically reduce the free model parameters. On the other hand approaches such as DeepChrome [22] and AttentiveChrome [23] reduce the inputs by binning them at 100 bases by bin.

Our proposed architecture was designed with the following needs in mind: first, to drastically reduce the parameter of the model; second, to provide a large receptive field; third, to minimize the meta-parameter search; and finally to minimize the introduction of implicit assumptions about biological phenomena.

The proposed architecture consists of three stages similar to the architecture described by Kim et. al [28], see Fig. 2. The first stage consists of a 1×1 convolution, which maps the histone channels to the internal channel representation. In the second stage, a block of layers with N input and N output channels is applied recursively as often as the (theoretical) receptive field we want to achieve. The third stage collates the outputs from each application of the second stage to a multiscale representation, pools across it, and feeds it into a fully connected layer followed by the output layer. N is the only metaparameter of the proposed architecture and can be perceived as the network’s capacity. The second stage recursive-block consists of a cascade of an inception-like layer ([29]), a ReLU non-linearity, and a batch normalization ([31]) layer; thus the output-features have the same size as the input-features and follow a normal distribution. The inception-like layer consists of N/2 1×1 convolutions and N/2 3×1 convolutions, padded as needed to ensure the output having the same size as the input. Each application of the recursive-block is followed by a striding of 2 allowing for an exponentially growing receptive field (RF) given by the formula:

**Fig. 2.**
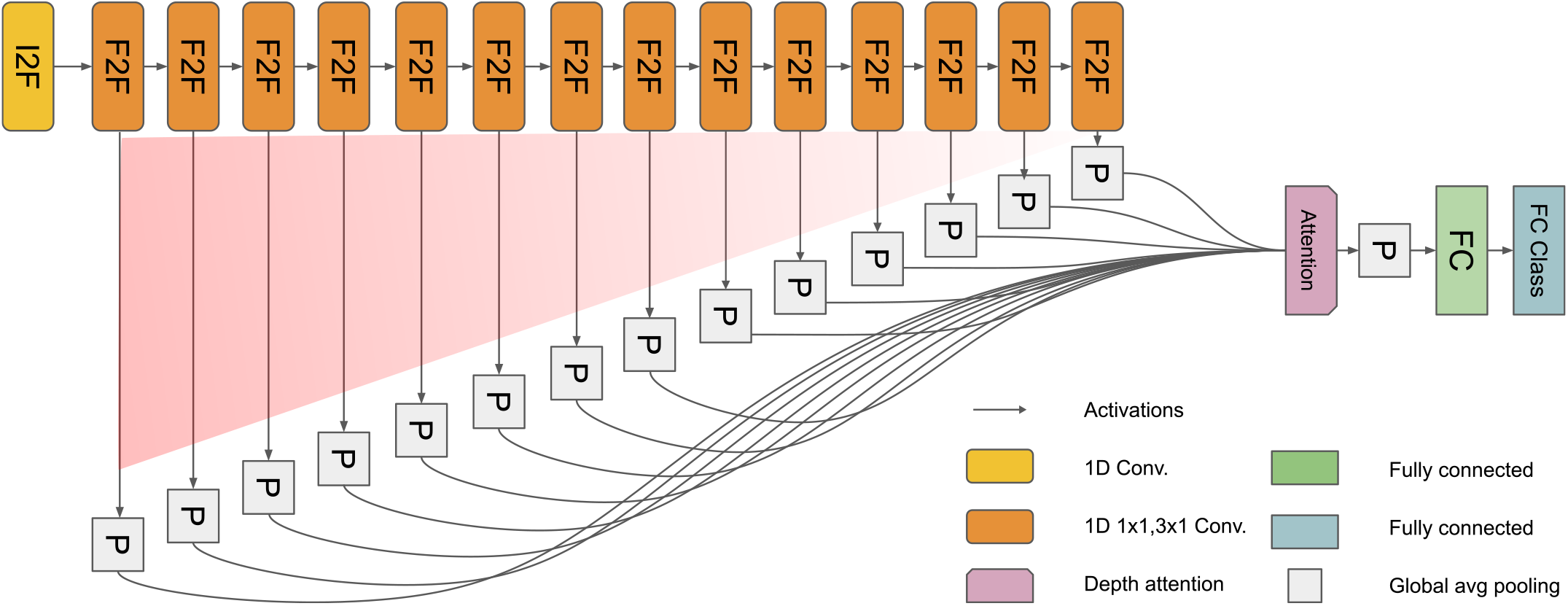
Model architecture consisting of one convolutional layer followed by a recursive block and a reception-like layer, “feeding” an attention-like mechanism, followed by two fully connected layers. Layers with shared weights have the same color. The activation’s spatial resolution is halved after each F2F layer.

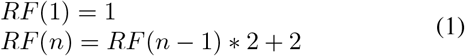

In the proposed architecture, the depth was fixed at a maximum of n=14 so that the receptive field of the ultimate convolutional layer is 24,574. The design that allows an exponentially growing receptive field, is inspired by Wavenet [32]. In Fig. 2 orange boxes represent the successive applications of the recursive block. The output of each application of the recursive-block are pooled into a single feature vector representing a particular scale. In the third stage these vectors are weighted by an attention-like mechanism and pooled into a single vector representing the full signal which is finally classified by the fully connected layers.

Fig. 3 illustrates that the first application of the recursive layer is emphasized, the following early applications contribute less and then they converge to one. As can be seen in Fig. 3 inference is realised through many alternative paths and the recursive-block can be perceived as the router.

**Fig. 3.**
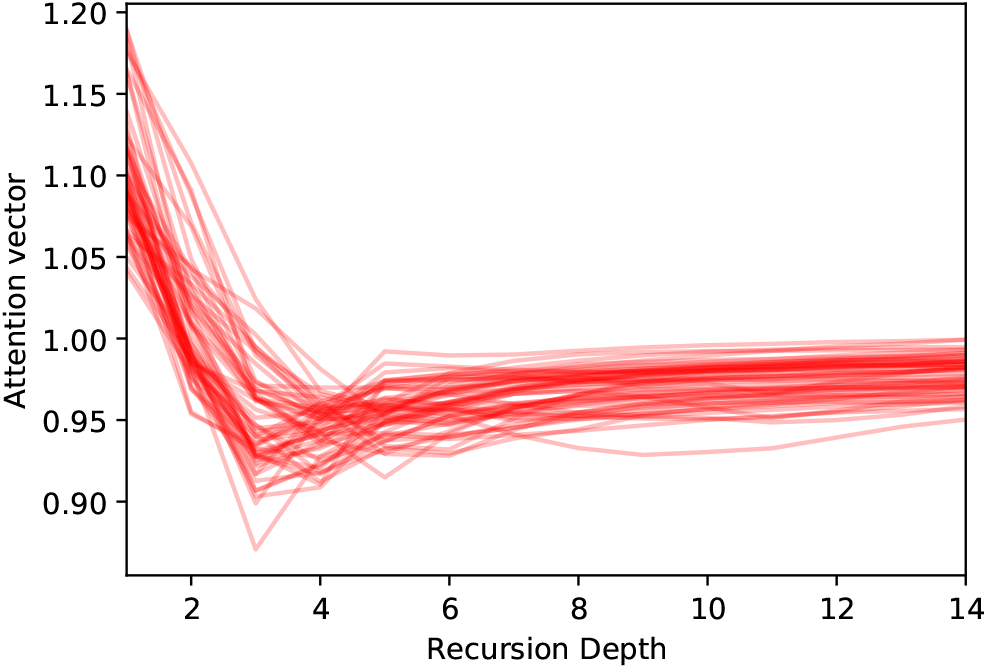
Attention-like vectors as a function of recursion depth based on the proposed architecture, trained for all 56 human cell lines (each line corresponds to the attention of a given cell line). First application of recursive layer is highly important; all applications converge to 1.

## IV. Experiments

We performed experiments in order to compare the proposed networks to the state-of-the-art and also in order to make informed decisions during the design of the proposed method. We limited the use of the test-sets to the proposed models and the state-of-the-art models. The validation set was used for the determination of the network architecture and its parameters. The test-set was only used after we had committed to the proposed networks in order to evaluate them. For comparing our proposed architecture with the state-of-the-art, we implemented the network described in DeepChrome ([22]) and trained, validated and tested it with our dataset. In addition, we used the publicly available model and train of AttentiveChrome ([23]) and tested it on our dataset, as described in IV-B.

### A. Dataset

In order to obtain uniform data of a large collection of chromatin marks, as well as quantification data of RNA expression in a variety of different samples, we used the REMC database ([9]). From the REMC database we obtained data for five core histone modification marks (H3K4me1/3, H3K36me3, H3K27me3 and H3K9me3) in 56 cell lines and the corresponding normalized RPKM ([33]) expression data for these cell lines. We selected these histone marks not only because of their availability, but also because they are implicated in the gene expression regulation, acting either as promoters, suppressors, or mediators ([4], [34], [35], [7]), see Table I. To place the genes on the hg19 genome we used an updated annotation of the genes, the Ensembl GRCh37 version 87 ([36]), in order to have the data reflect a current state of gene annotation, removing the expression data for the genes that were not included in the version 87 annotation. During this process approximately 100 genes were removed from the initial dataset. For each cell line, we partitioned 19796 genes as listed by [9] of the human genome into three equal sets for training, validation, and testing process, making sure that the three sets were completely isolated from each other. Genomic sequences that overlapped, either because they are located on opposite strands or overlap within the same genomic context, i.e. area around Transcription Start Site (TSS), were placed in the same set, in order to avoid samples in different partitions being near. We specifically guarantied a distance of 100,000 bases between any pair of samples coming from different partitions ([37]). The test-set was only used for comparing our model to the state of the art, while we used the validation set for any other experiment, tuning etc. For the labeling of the genes in the two categories of expressed or not-expressed, in each cell line, the gene expression values were discretized using the median normalized expression value of all genes in the cell line, in accordance with ([22], [23]). Genomic elements, such as genes and histone modifications, have a set of coordinates containing the chromosome on which the element is located and the starting and ending coordinates of the element on the given chromosome. During pre-processing, genomic coordinates are transformed into a uniform continuous space, transferring the information of the location into a two-element structure. This strategy can also be applied to different organisms with varying number of chromosomes and varying chromosome lengths. For each gene in the dataset, we focused on a genome context of maximum 15,000 bases upstream and downstream of the Transcription Start Site (TSS), thus remaining within the range of reported short range interactions between histone modifications and gene expression ([38], [39]). Within our implementation the histone signal is calculated in bins of size 1 base or higher, without restrictions on the size of the bins. In accordance to previous studies ([22], [23]), we made extensive experiments using a genome context of 5 000 bases upstream and downstream of the TSS and a bin size of 100 bases, although we also made experiments with higher genome context and variable bin sizes.

**TABLE I.**
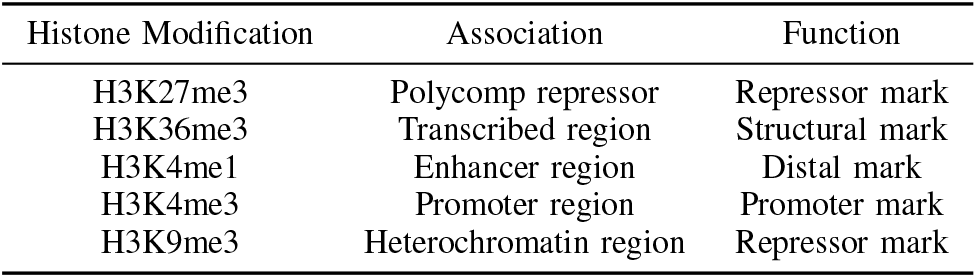
Histone modification marks and their functionality.

| Histone Modification | Association | Function |
| --- | --- | --- |
| H3K27me3 | Polycomb repressor | Repressor mark |
| H3K36me3 | Transcribed region | Structural mark |
| H3K4me1 | Enhancer region | Distal mark |
| H3K4me3 | Promoter region | Promoter mark |
| H3K9me3 | Heterochromatin region | Repressor mark |

### B. Baselines

#### a) DeepChrome

This classical CNN architecture is designed specifically for the task of chromatine sequence classification ([22]). For the experiments, we used our implementation of the exact model described in DeepChrome ([22]) and trained it with the given training parameters, ie a learning rate of 0.001, for 100 epochs, dropout of 0.5 was applied on the convolutional layer. DeepChrome receives inputs of 100 values for 5 channels. Each input value has the average histone signal for 100 bases so it covers 5 000 bases upstream and 5 000 bases downstream from the TSS. As the model has no global-pooling it can only operate on inputs of a fixed length of 100 bins.

#### b) AttentiveChrome

A Recurent Neural Network with two attention mechanisms that was proposed as a means of improving interpretabillity through the attention mechanism. Performance-wise it is inconclusive whether AttentiveChrome or DeepChrome performs better. We incorporated the publicly available implementation into our experimental pipeline; specifically we used the model definition and the training source code as they were and we simply adapted our data-loading and evaluation code to use the model. We used the exact training protocol of the public code. For both DeepChrome and AttentiveChrome, the validation and test set AUC reported is the best achieved during training. Although this is favorable to larger models that suffer from over-fitting, it can be argued that it is equivalent to early stopping. It can be inferred from the public source code that this is the employed evaluation protocol. The inputs to the network are the same as in DeepChrome.

### C. Thin, Slim, and Starved networks

The proposed architecture is designed with the goal of limiting the meta-parameter search. We did not observe any effect on performance when the recursion depth increased. We used a depth of 14 as we wanted to achieve a receptive field greater than 10 000 base-pairs. The only meta-parameter the proposed model has is the channel-count *N* which functions as a quantification of the model’s capacity. The parameter count of the proposed model is *iN* + 3*N* ^2^ + 5*N* + 2 where *i* is the number of input channels, which is equivalent to *O*(*N* ^2^) as *i* can be considered a constant. In our data it was 5. Three variants of the network are proposed: a thin variant that has 100 channels resulting in 31 016 parameters, a slim variant having 30 channels resulting in 3 076 parameters, and a starved variant that has only 10 channels resulting in 416 parameters. The fact that our networks only vary in capacity, allows for experiments on the capacity-performance trade-off which is a way to quantify the hardness of the problem.

### D. Comparison with the state-of-the-art

The proposed ReChrome network performs better than both DeepChrome and AttentiveChrome over the test-set as can be seen in table II. For every cell line of the 56 we used in experiments, networks were trained four times on the train genes, used the validation genes for early stopping and evaluated the ROC-AUC. This allows to have an estimate on the significance of performance measurements for each cell line independently. In Fig. 4 each model’s performance in each cell line can be seen; the standard deviation for the four replicates is marked as a black error range in the bars. ReChrome variants demonstrate smaller deviation than AttentiveChrome, while DeepChrome demonstrates to be more volatile. As there are 56 cell lines, each one being a dataset for gene expression in a different biological context, statistical testing can allow for safe estimates on the performance of the proposed ReChrome and the state-of-the-art (DeepChrome and AttentiveChrome). All ReChrome variants, ReChrome, ReChrome-Slim, and ReChrome-Starved outperform DeepChrome with p-values 1.63E-45, 1.43E-46 and 3.12E-42 respectively. ReChrome and ReChrome-Slim outperform AttentiveChrome as well, with p-values 2.67E-05 and 8.83E-06 respectively. AttentiveChrome outperforms ReChrome-Starved variant, although not significantly (p-value 0.007). While DeepChrome and AttentiveChrome were reported to have similar performance by their authors, we find that AttentiveChrome is more consistent, robust, and less susceptible to over-fitting. The early-stopping algorithm we used, was to train each model for 20 epochs and keep the model at the point were it reached it’s top performance on the validation set. While this algorithm is good for well regularised models such as ReChrome and AttentiveChrome, if a model is not regularised such as DeepChrome, this algorithm might lead to overfitting the validation-set.

**Fig. 4.**
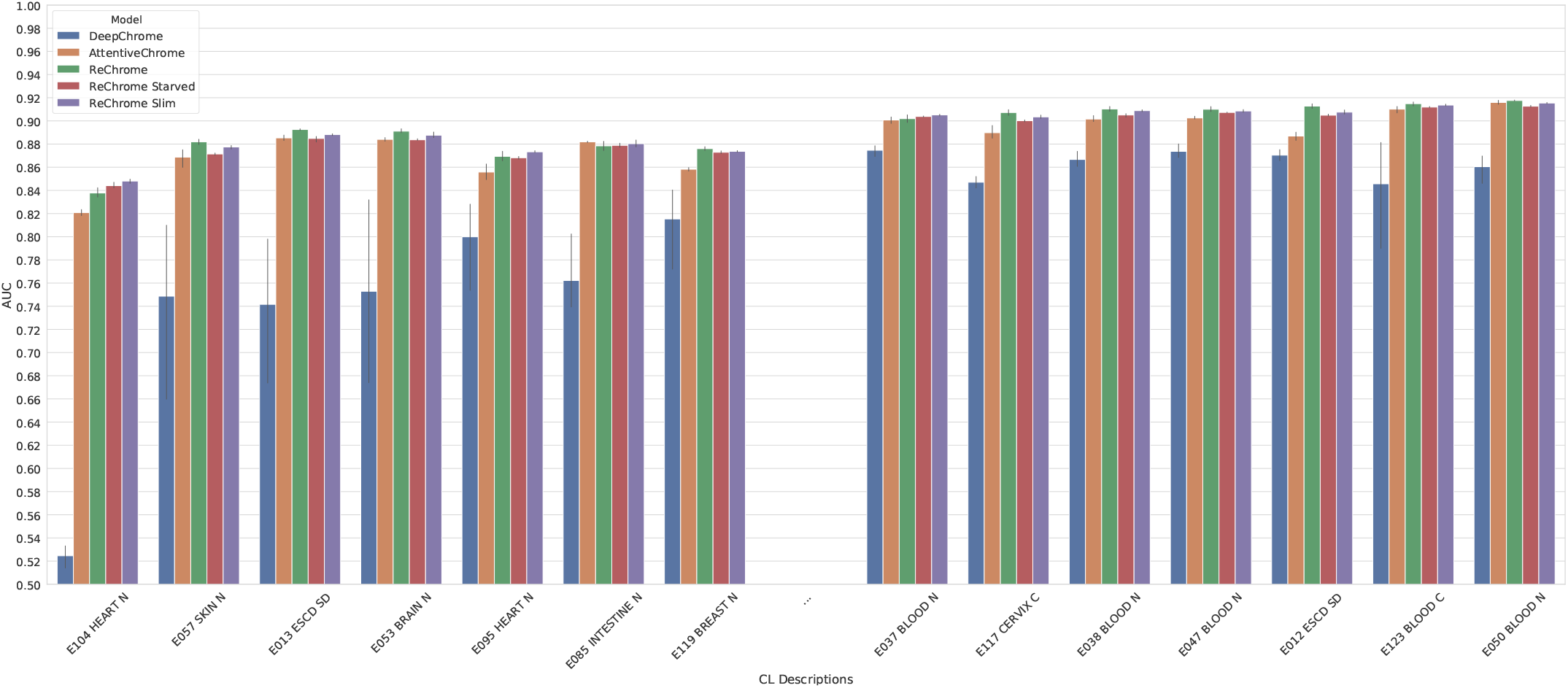
Performance of the different variants of ReChrome compared with the state-of-the-art, on the 7 cell lines with the lowest performance and the 7 cell lines with the highest. The thin variant is performing better than the slim and starved.

### E. Cross Dataset Generalization

In order to assess how well the networks generalize, we conducted experiments where each network is trained in one cell line and then tested in all remaining 55 cell-lines. This experiment provides intuitions on the importance of biological context for determining the relationship between chromatin marks and gene-expression. As can be seen in the last column of Table II, on average performance of Rechrome and ReChrome-Slim trained in one cell-line and tested on an other only dropped by 1.6% and 1.2% respectively while ReChrome-Starved had no observable performance drop at all. These findings validate our hypothesis that a drastic reduction in the neural networks capacity, will force the models to learn more generic associations. ReChrome-Starved, has 416 parameters and was trained on 6598 samples of size 5 × 100, and manages to generalise perfectly between different biological contexts.

### F. Versatility

A benefit of the recursive architecture of ReChrome, is that it allows the exact same model to be employed on data of arbitrary size. We performed several experiments by altering the sampling parameters of ReChrome’s input. In Table II all networks had 100 inputs per channel each input bin had the average of 100 bases proving a context of ±5,000 bases around the TSS of each gene. In Table III the average performance of ReChrome on the validation-set of each cell-line with alternative sampling parameters can be seen. We tried bin sizes of 1, 100, 150, 300, 400 and 15,000 bases for contexts of approximately 10,000 and 30,000 bases. It can be seen that on several occasions ReChrome benefits and even improves it’s performance on validation set compared to the standard sampling parameters.

**TABLE II.** Average AUC of the different models over validation, test and cross-test set.

| Model | Parameters | Val. set | Test set | Cross test set |
| --- | --- | --- | --- | --- |
| DeepChrom | 644,177 | 87.36% | 81.43% | 79.78 |
| AttentiveChrom | 55,681 | 86.36% | 86.8% | NA |
| ReChrom | 31,016 | **87.54%** | 87.73% | 86.13% |
| ReChrom Slim | 3,076 | 87.06% | **87.75%** | **86.56%** |
| ReChrom Starved | 416 | 86.55% | 86.45% | 86.45% |

**TABLE III.** Average AUC of the different bin-sizes and bin-count on Validation.

| Model | bin size | bin count | TSS context (bases) | AUC % |
| --- | --- | --- | --- | --- |
| ReChrom | 1 | 30,000 | $\pm 15,000$ | 85.35 |
| ReChrom | 100 | 100 | $\pm 5,000$ | 87.54 |
| ReChrom | 150 | 66 | $\pm 4,950$ | 87.64 |
| ReChrom | 150 | 200 | $\pm 15,000$ | 87.63 |
| ReChrom | 300 | 34 | $\pm 10,200$ | 88.05 |
| ReChrom | 300 | 100 | $\pm 15,000$ | 88.07 |
| ReChrom | 400 | 26 | $\pm 5,200$ | **88.14** |
| ReChrom | 400 | 76 | $\pm 15,200$ | 88.09 |
| ReChrom | 15,000 | $\pm 2$ | 87.35 | |

### G. Visualization

In order to gain meaningful visualizations of what ReChrome actually learns, we employed gradient ascent to construct optimal synthetic inputs that maximize the network’s confidence for both expressed and non-expressed classes. In Fig. 5 we can see optimal inputs for both classes for the DeepChrome baseline and ReChrome. The gradient ascent was obtained with a learning rate of 0.01, weight decay coefficient *λ* 0.001 and 600 iterations. It is worth remarking that the proposed ReChrome has observed very clearly the relationship of the existence and absence of chromatin marks and gene expression. Presence of structural, distal and promoter marks, and absence of repressor marks is found to be the optimal input for expressed genes, while the exact opposite situation is found to be the optimal input for non-expressed genes. ReChrome is in agreement with known functional relationships between gene expression and specific histone modifications Table I, while it seems DeepChrome focuses on complicated interactions at the expense of the dominant linear ones. This observation demonstrates that ReChrome, while sensitive to non-linear variable interactions that allow it to outperform the state-of-the-art, tends to produce classifications that are also consistent with the fact that H3K4me1, H3K4me3 and H3K36me3 are known activators and H3K27me3 and H3K9me3 known supressors of gene expression. [40], [4], [7], [38], [14], [6], [41].

**Fig. 5.**
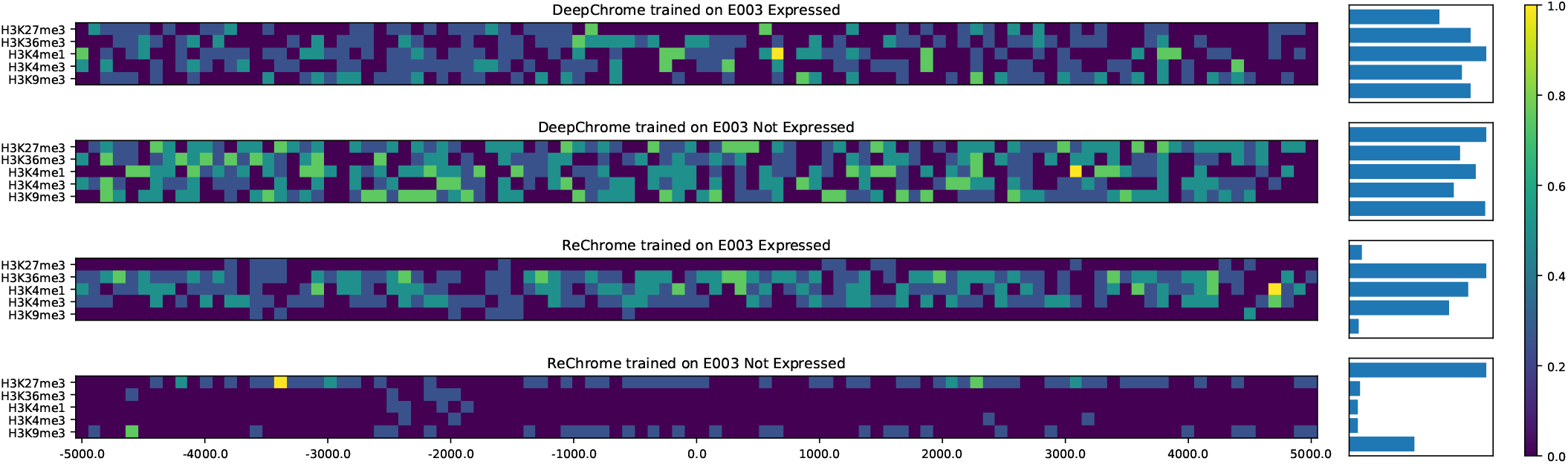
Gradient ascent visualization of the proposed ReChrome and DeepChrome models on both expressed and non-expressed samples of E003 (Embryonic Stem Cells).

## V. Conclusions

### A. Findings

In this work, we proposed a recursive convolutional neural network that drastically reduces the number of parameters and meta-parameters and at the same time, manages a small but significant improvement over the state-of-the-art. The proposed architecture is well regularised because of its constrained capacity and proved to be impervious to over-fitting of any kind. We consider that the value of predicting gene expression from histone marks does not lie in using machine learning to reduce the cost of analysis but rather, on providing a valid quantification and a lower bound of the interaction between these two phenomena.

The proposed architecture dealt very effectively with the presented data and proved to be highly stable, one might argue that this is because the proposed model can specifically discover the suppressor or promoter roles each histone mark has as seen in Fig. 5. We should point out that established linear and non-linear methods can not cope as well as deep learning [22] with this kind of data. Yet the scope of domains to which ReChrome can successfully be employed remains a question.

### B. Discussion

While the proposed model outperformed significantly the state-of-the-art; the error rate of all alternative architectures, is very large for a binary classifier and there might be a limit to the performance machine learning techniques can achieve with the specific data used in these experiments. Apart from the histone modifications that we have explored, several others exist that may also have an impact on gene expression. The addition of more modifications could also improve the prediction of gene expression. DNA methylation ([42]) is another layer of epigenetic regulation, which also impacts gene expression. It gives an additional layer of information and another channel that could be used as input to our model. We used a genomic context of 10,000 bases around each gene’s TSS; but there are histone marks that are modulating gene enhancers, some of which might be even millions of bases further away from the genes that they regulate [43], [40], [44]. In fact, a gene can have an accumulation of active marks and still be non-active, because the long range enhancer that participates in the regulation of the gene is not in an active state [45].

While these enhancers are far away from the genes if we regard the DNA as a 1D structure, this does not reflect the reality, because the DNA has a 3D structure [46], specifically it is a thread folded into knots several times over; even-more the knots fold and unfold dynamically and present day acquisition protocols average over different such states. Nonetheless, the addition of data regarding the structure of the DNA, such as Hi-C data [47], would be valuable.

### C. Future Work

The work presented in this paper, was principally aimed at establishing that ReChrome is a reliable tool for analysis of epigenomic data across different configurations. Our experiments were principally aimed at comparing fairly ReChrome to the state-of-the-art instead of aiming at biologically meaningful insights. The binary expressed, not expressed classes obtained by thresholding expression values to above and bellow median, leaves much to be desired. Three classes might make more sense as under-expressed, normal, over-expressed genes. These classes should not be defined by ranked order statistics as this forces assumptions on the population of gene expression. Although experiments in this paper demonstrated that ReChrome works and generalises across several different cell-types of humans, experiments for other species including plants and other mammals are planned.

## Acknowledgment

We gratefully acknowledge the support of NVIDIA Corporation with the donation of the Titan V GPU used for this research.

